# Protein-based Immunome Wide Association Studies (PIWAS) for the discovery of significant disease-associated antigens

**DOI:** 10.1101/2020.03.18.997759

**Authors:** Winston A. Haynes, Kathy Kamath, Patrick S. Daugherty, John C. Shon

## Abstract

Identification of the antigens associated with antibodies is vital to understanding immune responses in the context of infection, autoimmunity, and cancer. Discovering antigens at a proteome scale could enable broader identification of antigens that are responsible for generating an immune response or driving a disease state. Although targeted tests for known antigens can be straightforward, discovering antigens at a proteome scale using protein and peptide arrays is time consuming and expensive. We leverage Serum Epitope Repertoire Analysis (SERA), an assay based on a random bacterial display peptide library coupled with NGS, to power the development of Protein-based Immunome Wide Association Study (PIWAS). PIWAS uses proteome-based signals to discover candidate antibody-antigen epitopes that are significantly elevated in a subset of cases compared to controls. After demonstrating statistical power relative to the magnitude and prevalence of effect in synthetic data, we apply PIWAS to systemic lupus erythematosus (SLE, n=31) and observe known autoantigens, Smith and Ribosomal P, within the 22 highest scoring candidate protein antigens across the entire human proteome. We validate the magnitude and location of the SLE specific signal against the Smith family of proteins using a cohort of patients who are positive by predicate anti-Sm tests. Collectively, these results suggest that PIWAS provides a powerful new tool to discover disease-associated serological antigens within any known proteome.

**Author Summary:** Infection, autoimmunity, and cancer frequently induce an antibody response in patients with disease. Identifying the protein antigens that are involved in the antibody response can aid in the development of diagnostics, biomarkers, and therapeutics. To enable high-throughput antigen discovery, we present PIWAS, which leverages the SERA technology to identify antigens at a proteome- and cohort-scale. We demonstrate the ability of PIWAS to identify known autoantigens in SLE. PIWAS represents a major step forward in the ability to discover protein antigens at a proteome scale.

## Introduction

Antibodies present in human specimens serve as the primary analyte and disease biomarker for a broad group of infectious (bacterial, viral, fungal, and parasitic) and autoimmune diseases. As such, hundreds of distinct antibody detecting immunoassays have been developed to diagnose human disease using blood derived specimens. The development of high-throughput sequencing technologies has enabled sequencing of numerous proteomes from diverse organisms. However, methods for antigen discovery within any given proteome remain relatively low throughput. The serological analysis of expression cDNA libraries (SEREX) method has been applied frequently to identify a variety of antigens, but high quality cDNA library construction remains technically challenging and time consuming [1–3]. Alternatively, entire human and pathogen derived proteomes can be segmented into overlapping peptides, and displayed on phage or solid-phase arrays and probed with serum [4–6]. Fully random peptide arrays of up to 300,000 unique sequences have also been used successfully to detect antibodies towards a range of organisms [7–9]. Even so, the limited molecular diversity of array based libraries can reduce antibody detection sensitivity and hinder successful mapping of petide motifs to specific proteome antigens [7]. Thus, a general, scalable approach to identify serological antigens within arbitrary proteomes is needed.

In autoimmune diseases and cancers, autoantigen discovery is further complicated by the size of the proteome, heterogeneity of disease, and variability in immune response. Patient genetics, exposures, and microbiomes contribute to this heterogeneity, which in turn yields disparate responses to diverse antigens and epitopes [10,11]. In such cases, the mapping of multiple epitopes to one antigen can increase confidence in a candidate antigen [7,12]. Even for diseases with conserved autoantigens, epitope spreading can lead to a diversified immune response against additional epitopes from the same protein or other proteins from the same tissue [13,14]. In cancer patients, neoepitopes can arise in response to somatic mutations that yield conformational changes or abnormal expression [15,16].

In complex autoimmune diseases like systemic lupus erythematosus (SLE), autoantibodies play an important role in diagnosis, patient stratification, and pathogenesis. SLE autoantigens include double-stranded DNA, ribonuclear proteins (Smith), C1q, α-actinin, α-enolase, annexin II, annexin AI, and ribosomal protein P [17–19]. In particular, anti-Smith antigen antibodies are present in 25-30% of SLE patients [20,21]. The Smith antigen consists of a complex of U-rich RNA U1, U2, U4/U6, and U5, along with core polypeptides B’, B, D1, D2, D3, E, F, and G. Not all components of this complex are equally antigenic, and there are multiple epitopes within the complex [22,23].

One approach for antigen discovery, serum epitope repertoire analysis (SERA), uses bacterial display technology to present random 12mer peptides to serum antibodies [24–26]. Peptides that bind to serum antibodies are separated using magnetic beads and sequenced using next-generation sequencing. For each of these peptides and their kmer subsequences, enrichment can be calculated by comparing the actual number of observations to that expected based on amino acid frequencies [24]. Mapping these peptide epitopes to their corresponding protein antigens requires protein structure and/or sequence. Structure-based epitope mapping methods (e.g., 3DEX, MIMOX, MIMOP, Pepitope) are not yet feasible at a proteome scale, due in part to the large number of undetermined structures [27–30]. However, since 85% of epitope-paratope interactions in crystal structures have a linear stretch of 5 amino acids, sequence information alone can be sufficient to identify many antigens [31–33]. The K-TOPE (Kmer-Tiling of Protein Epitopes) method has demonstrated the ability of tiled 5-mers to identify known epitopes in a variety of infections at proteome scale [34]. Here, we present a method, Protein-based Immunome Wide Association Study (PIWAS), which leverages the SERA assay to discover disease relevant antigens within large cohorts and at proteome scale. We evaluate PIWAS with synthetic data to examine the magnitude and prevalence of the effect needed for robust detection. We validate PIWAS using specimens from individuals with SLE and controls, identifying established anti-Smith and anti-Ribosomal P autoantibodies. We further validate the anti-Smith epitopes identified in our analysis using specimens positive for anti-Smith autoantibodies by predicate tests.

## Results

### PIWAS allows identification of proteome-based signals

To identify candidate serological antigens from arbitrary proteomes, we developed a robust, cohort-based statistical method to analyze peptide sequence data from the SERA assay. SERA uses a large bacterial display random peptide library of 10 billion member 12mers to identify binding to the epitopes recognized by antibodies species in a biospecimen (e.g. serum, plasma, cerebrospinal fluid) [Figure 1A]. From a typical specimen, we acquire 1-5 million unique 12mers. We break these 12mers into their constitutent kmers, calculate log-enrichments (observed divided by expected counts), and store the results in a BigTable database. To identify disease-specific antigens from these data, PIWAS compares kmer data from case and control cohorts against a proteome of interest (Figure 1B). For each protein and specimen dataset, we calculate tiled kmer enrichments (normalized to the controls as a background) and smooth across a sliding window. For each protein, we leverage statistics such as the outlier sum and Mann-Whitney test to compare the case and control populations. At a proteome scale, we prioritize candidate antigens based on these statistics (see Methods).

**Figure 1.**
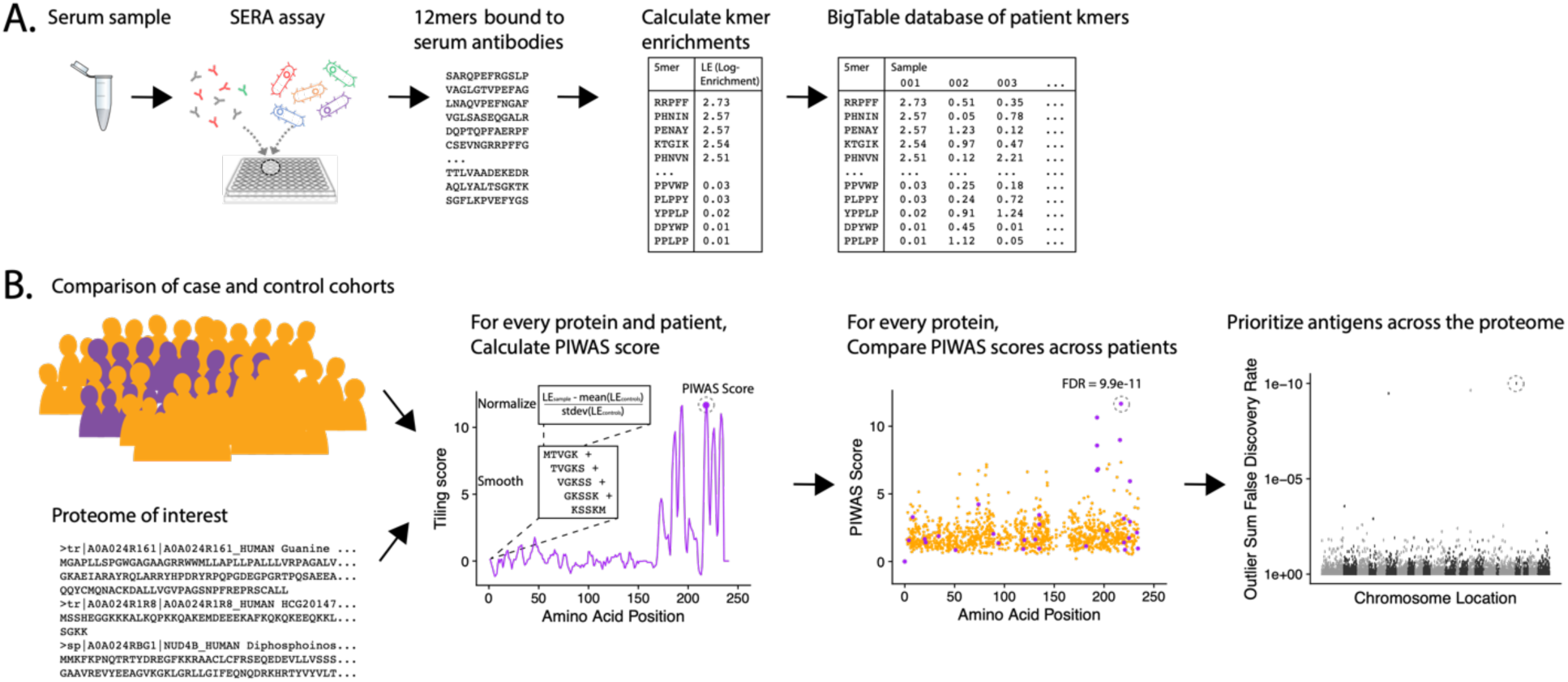
PIWAS discovers candidate disease antigens through proteome-wide analysis. (A) Case and control specimens are processed using SERA to generate a dataset of 12mer amino acid sequences bound by serum antibodies. Each 12mer is broken into kmer components and log-enrichments of these kmers are calculated, where enrichment indicates the number of observations compared to expectation based on amino acid frequency. (B) As input for the PIWAS algorithm, case and control cohorts are identified (purple, cases; gold, controls) as well as the target proteome. For each individual in the case and control cohorts and protein in the proteome, PIWAS scores are calculated by tiling kmers onto the protein sequence, smoothing over a window of these kmers, normalizing to the background signal in the controls, and calculating the maximum value. PIWAS scores are compared across all case and control samples to detect proteins whose scores are significantly greater in some subset of the case population than in the control population. Antigens are then rank-ordered by one or more statistics across the entire proteome.

### Kmer enrichment in samples with serum compared to enrichment in a random library

We first compared SERA library sequence composition before and after library selection with serum from healthy controls and SLE patients (Figure 2 A,B). Both the control and SLE serum yielded larger enrichments for both 5mers and 6mers compared to the unselected library. The enrichment of 5mers and 6mers in samples incubated with serum demonstrates the effects of antibody selection on the peptide library composition. We also compared the distribution of PIWAS values when 5mers were mapped to the human proteome. Interestingly, both SLE and anti-Smith cohorts yielded PIWAS value distributions with longer tails when analyzed against the entire human proteome when compared to those of healthy controls (Figures 2C). These findings confirm the general basis for using 5mers and 6mers for identifying both enriched signal in serum relative to a random library and enriched autoantigen signal using PIWAS in an example disease population relative to healthy controls.

**Figure 2.**
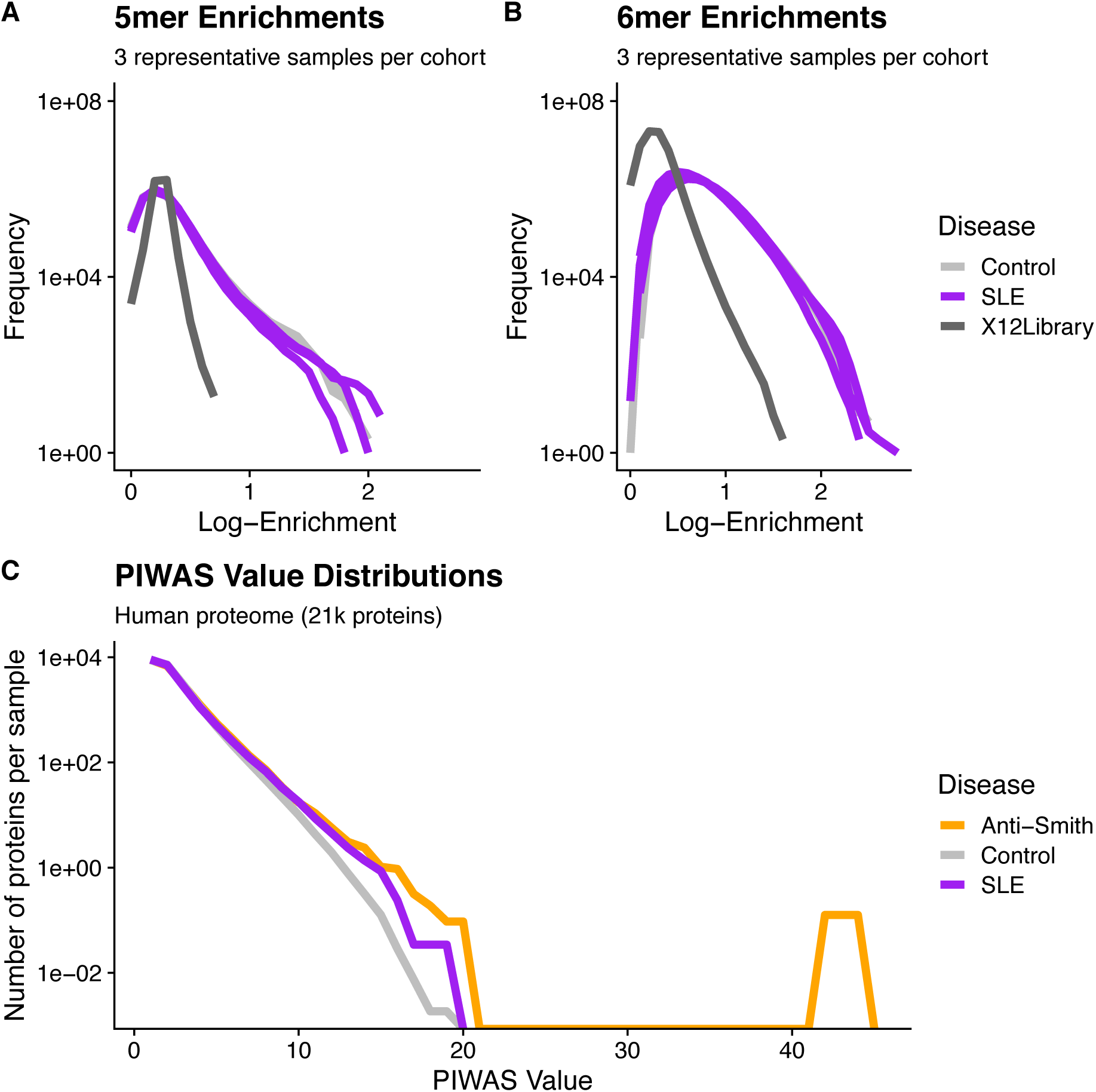
Distributional differences in kmer enrichments and PIWAS values between the unselected library and after selecting with SLE and control specimens. 5mer (A) and 6mer (B) Kmer frequency (y-axis) vs. Log-enrichment score (x-axis) for 6 subjects and the naïve library demonstrates species with large enrichments are found exclusively in those SERA assays incubated with serum. All 5mers or 6mers from three representative samples per cohort are evaluated for enrichment. Dark-gray lines = naïve 12-mer peptide library, purple lines = SLE cohort, gray lines = control cohort. (C) A comparison of PIWAS values (x-axis) vs. the number of proteins per sample with the corresponding PIWAS value (y-axis) reveals differences in both the range and distribution of PIWAS values between SLE and control samples. Distributions are based on 31 SLE cases and 1,157 controls. Purple = SLE cohort, gray = control cohort, orange = anti-Smith cohort.

### PIWAS Power Simulations

In order to assess the statistical power of PIWAS to detect enriched antigens in a cohort, we performed computational experiments where we adjusted the magnitude and prevalence of known autoantigenic signal against Sm antigens (specifically small nuclear ribonucleoprotein-associated proteins B and B’) in a cohort of SLE patients. Unsurprisingly, as the magnitude of the effect increases, so does the significance of the antigenic signal (Figure 3A). At an effect of only 60% of the SERA signal obtained with true SLE biospecimens, Sm antigens are significant at FDR=0.017 using the outlier sum FDR, still ranking within the top 20 proteins. Similarly, as the prevalence of the anti-Sm signal increases in the case population, so too does the significance of the outlier sum p-value (Figure 3B). At a prevalence of 7% (less than half of the actual biological prevalence in this cohort), anti-Sm is significant at FDR= 0.015 and remains within the top 20 scoring proteins.These results indicate an ability to detect signals well below the prevalence of many established autoantigens.

**Figure 3.**
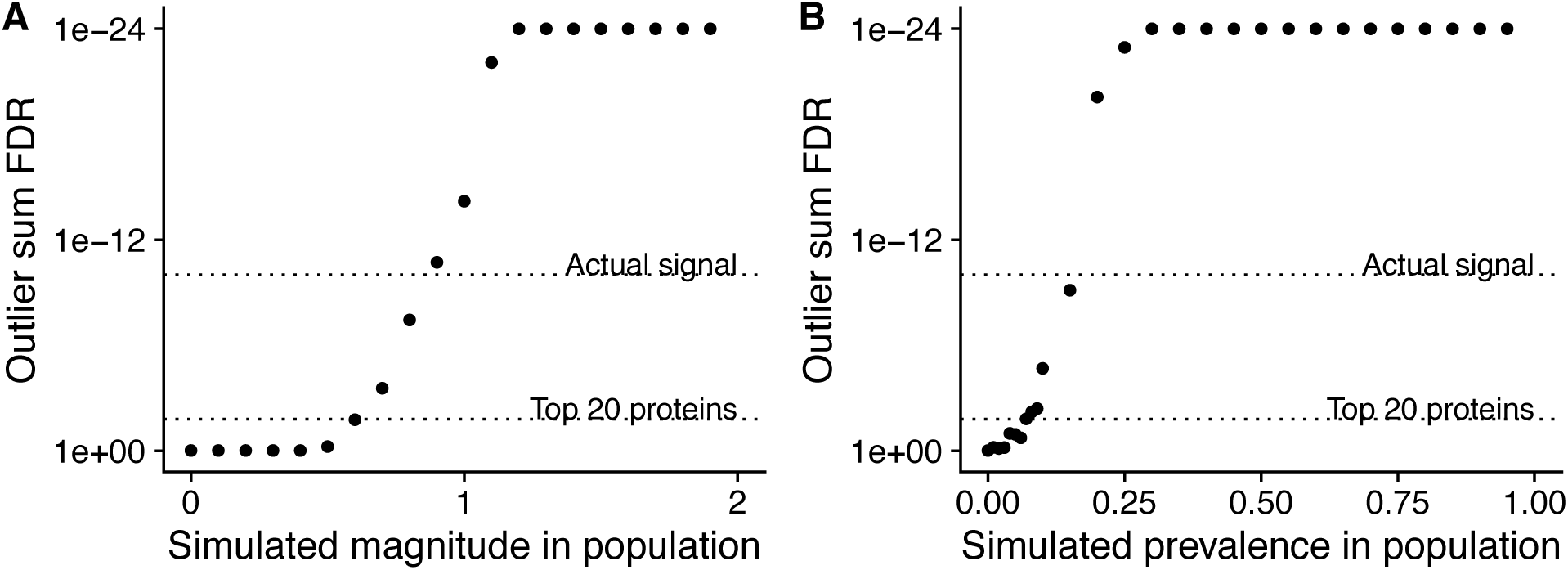
Simulations of magnitude and prevalence of autoantigenic signal to assess statistical limits of detection for PIWAS. SERA datasets from a cohort of SLE patients and kmer enrichments on small nuclear ribonucleoprotein-associated proteins B and B’ were used as the actual biological signal (magnitude = 1 and prevalence = 19%). The magnitude (A) and prevalence (B) of the kmer signal in this cohort was synthetically modulated to understand the statistical limits of detection for PIWAS.

### PIWAS analysis of SERA datasets from SLE specimens

We performed PIWAS to identify candidate autoantigens using specimens obtained from SLE patients. PIWAS results from individuals with SLE (n=31) were compared to those from controls (n=1,157) and proteins were ranked based on outlier sum FDR as a measure of significance across the human proteome (21,057 proteins) (Figure 4A-B). The highest scoring 22 proteins had outlier sum FDRs ranging from 1.6e-2 to 9.9e-11 and included multiple established autoantigens. Four Smith complex antigens were among the top seven hits with small nuclear ribonucleoprotein-associated proteins B and B’ exhibiting the highest significance (outlier sum FDR = 9.9e-11). In addition, 60S acidic ribosomal protein P1, another known SLE autoantigen [20,35], was highly significant. Multiple highly significant epitopes were evident within nuclear ribonucleoprotein-associated proteins B and B’ (Figure 4C, Table 1). The most significant enrichments occurred at two different locations near the C-terminus.

**Table 1.**
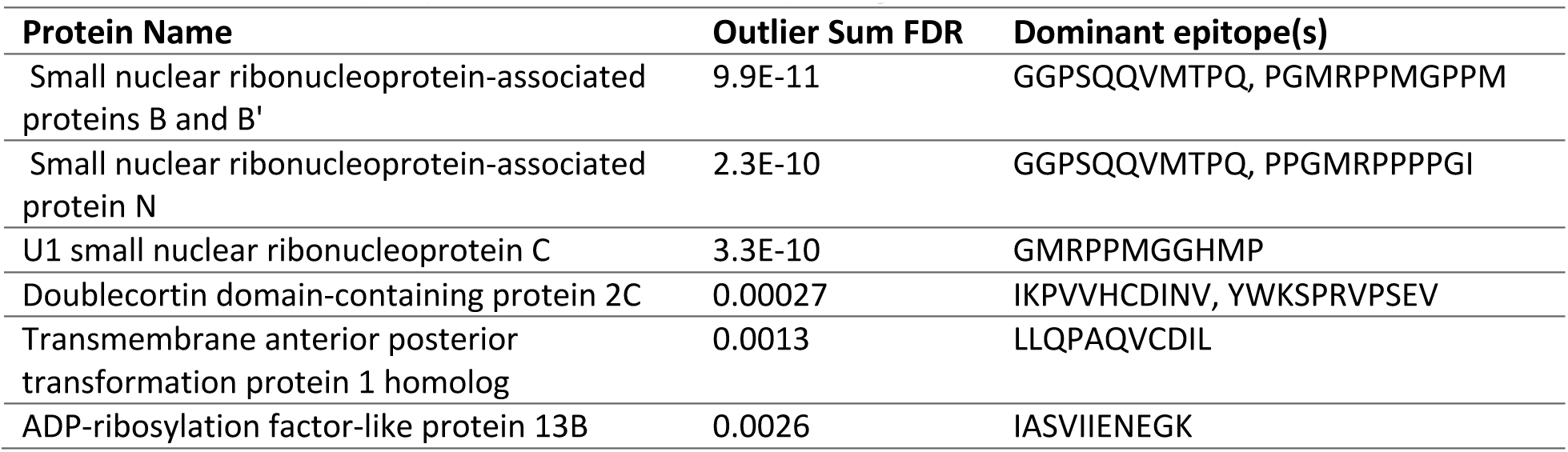
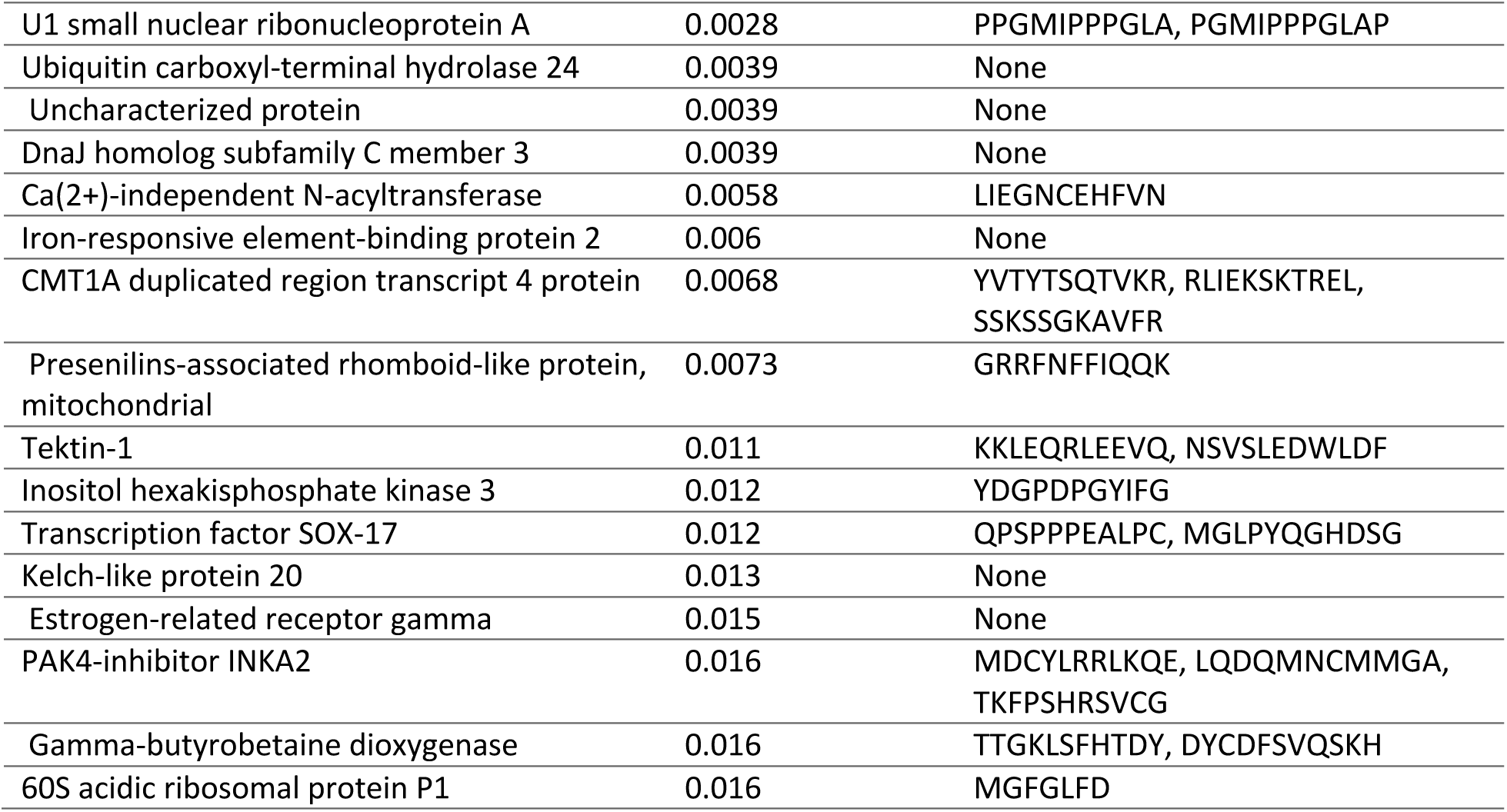
Dominant epitopes for highest scoring antigens from SLE PIWAS.

**Figure 4.**
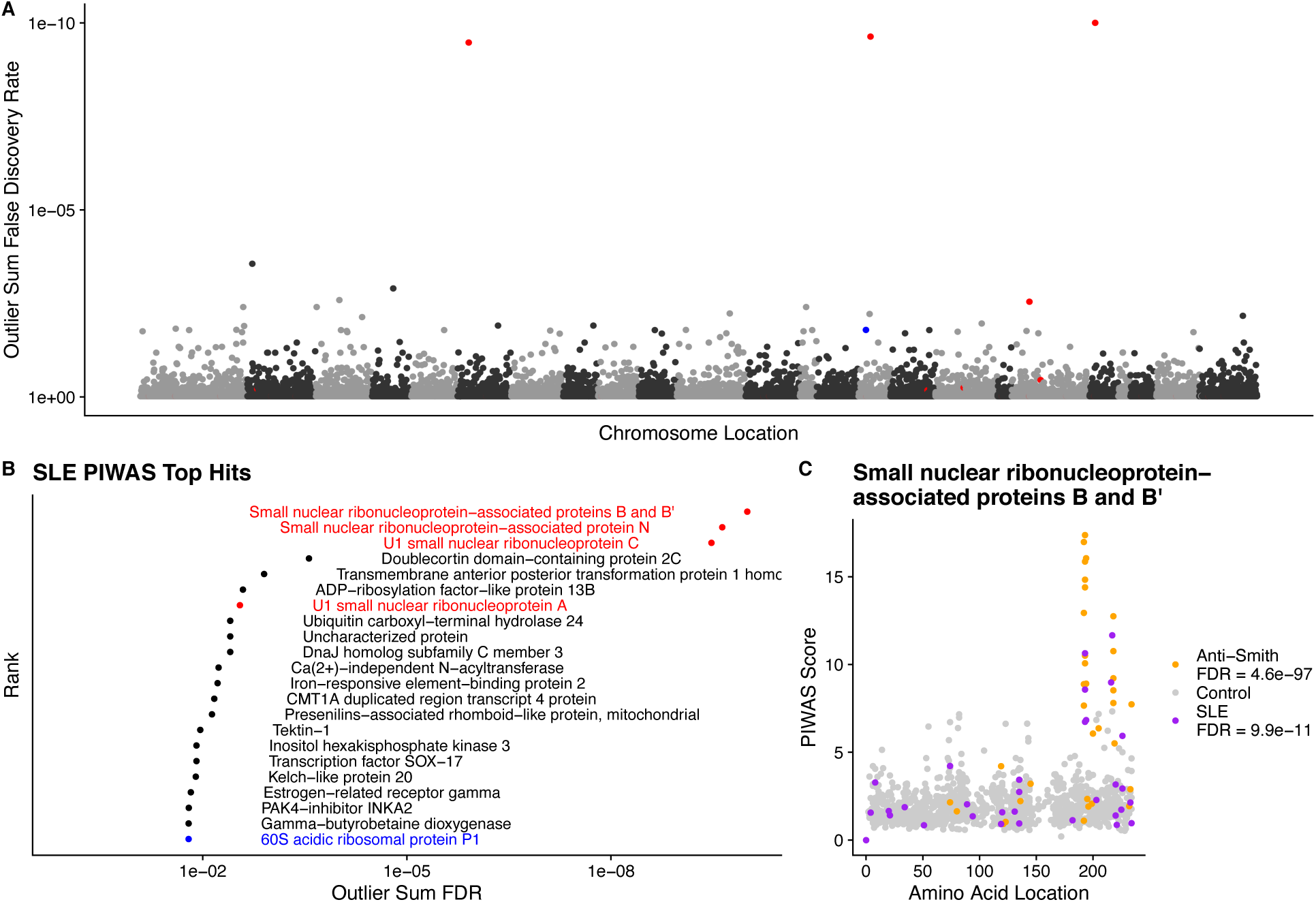
Literature reported and putative autoantigens are detected in SLE samples by PIWAS. (A) PIWAS results from a comparison of SLE samples to controls against the human proteome were prioritized using outlier sum FDR as a measure of significance (y-axis, see Methods). For visualization, proteins were laid out according to chromosome location. (B) Among the top set of 22 ranked proteins, 5 are established autoantigens (Smith family in red, others in blue). (C) Strength (y) and location (x) of PIWAS scores for the small nuclear ribonucleoprotein-associated proteins B and B’ within SLE (n=31, purple) vs. control (n=1,157, grey). A cohort of anti-Sm predicate positive patients (n=35, orange) were compared to the same controls to validate the signal obtained using SLE specimens with unknown anti-Sm serostatus.

### PIWAS in an independent cohort of Smith antigen positive subjects

To investigate the ability of PIWAS to identify Smith antigens in an independent cohort positive for anti-Sm using validated clinical tests, we applied PIWAS to a cohort of 35 Smith antigen positive samples. In this anti-Sm seropositive cohort, PIWAS again clearly identifies Smith antigens at the top of the ranked list of antigens (Table 2). The dominant C-terminal, anti-Sm epitope was identical between the two independent cohorts.The statistical significance within the second cohort is greatly increased relative to the general SLE cohort as might be expected, given the 100% seroprevalence of anti-Smith within this second specimen set. The unbiased identification of known SLE autoantigens in independent cohorts validates the ability of PIWAS to identify shared autoantigens in a data-driven way.

**Table 2.** Dominant PIWAS epitopes for top antigens from anti-Smith seropositive specimens.

| Protein Name | Outlier Sum FDR | Dominant epitope(s) |
| --- | --- | --- |
| Small nuclear ribonucleoprotein-associated protein N | 1.1e-98 | PGMRPPPPGIR |
| Small nuclear ribonucleoprotein-associated proteins B and B' | 4.6e-97 | PGMRPPMGPPM |
| U1 small nuclear ribonucleoprotein A | 1.2e-67 | PPGMIPPPGLA |
| U1 small nuclear ribonucleoprotein C | 1.3e-47 | PGMMPVGPAPG |

## Discussion

We demonstrate the utility of a general and scalable methodology to identify serological antigens within arbitrary proteomes using Protein-based Immunome Wide Association Studies (PIWAS). The power of PIWAS derives from cohort-based statistical analyses within large datasets of antibody-binding epitopes. PIWAS analyzes the enrichments of proteome spanning overlapping 5mers and 6mers that are observed amongst a peptide library selected for binding to antibody repertoires from cases and controls. We show that the kmer enrichment space demonstrates enriched signals compared to the unselected libraries. Further, the PIWAS space is enriched in SLE patients compared to control samples. Using synthetic data, we found that PIWAS has power to detect significant antigens at a signal of only 60% of the signal of a known autoantigen. When applied to experimental datasets from SLE cases and controls, PIWAS ranks SLE-specific Smith antigens highly in a proteome-wide search of candidate antigens. Finally, the epitopes from this antigen family were validated using a cohort of anti-Sm autoantibody positive patients.

Previous approaches to proteome-scale antigen identification rely on wet lab approaches that require a priori knowledge of the target proteome when the assay is performed [1–6]. In contrast, the use of random peptide library data with PIWAS enables analyses against arbitrary proteomes. In addition to the reference human proteome utilized here, the same SERA data can be reanalyzed against proteomes of infectious agents, patient-specific mutations, and splice variants, without performing additional wet lab assays. Indeed, we have identified previously validated epitopes for multiple bacterial, viral, and fungal infectious diseases using this method [data not shown].

PIWAS is an immunological analog to widely employed genome-wide association studies (GWAS) that employ statistical association of gene variants in large disease and control cohorts to identify disease-associated loci. Like GWAS, PIWAS employs a data-driven statistical approach to scan entire genomes and proteomes for statistically significant differences between case and control cohorts. Advancements in GWAS methods such as burden testing has enabled multiple variants witinin a single gene to be collapsed, thereby increasing the power to detect disease-associated genes [36,37]. Similarly, PIWAS scans each protein to find a maximum signal and allows for the contributions of multiple distinct epitopes to identify candidate antigens associated with disease. By leveraging the outlier sum statistic [38], we are able to further highlight antigens with signals that are strong, but present in only a subset of the patient population, or derive from unique epitopes within the same antigen.

Just as GWAS must consider a variety of biological and technical limitations, effective PIWAS must consider and address pre-assay, assay, and post-assay factors that can impact performance. The most significant pre-assay issues relate to the selection of cohorts for disease and control populations. Our analyses using synthetic data demonstrated that magnitude and prevalence of autoantigenic signal affects the ability of PIWAS to prioritize antigens. Thus, clean case and control cohorts are more likely to yield genuine autoantigens. In this study we were able to detect known antigens using a small cohort of SLE cases. As the cohort size grows, we anticipate even greater power to identify known and novel autoantigens.

Application of PIWAS to a cohort of SLE subjects identified known autoantigens, with 5 of 16 of the highest ranking hits across the entire human proteome being validated and clinically significant autoantigens. In particular, Smith antigens stood out as top hits in the SLE analysis. To validate this particular hit, we analyzed specimens from a second independent cohort of patients that tested positive for anti-Sm using clinical predicate tests. We found that the anti-Sm positive cohort exhibited reactivity against the same antigens and epitopes as the less homogeneous SLE discovery cohort. PIWAS identified an anti-Sm epitope ocurring within a proline rich region in agreement with multiple prior studies [20,39].

Other highly ranked proteins identified using PIWAS could represent novel candidate antigens associated with SLE. PIWAS ranks antigens based on the maximum signal observed across a cohort, however it is not always possible to determine which antigens are biologically significant due to sequence similarity between proteins. Therefore antigens ranked highly in PIWAS should be considered candidate antigens, and orthogonal experimental validation is generally necessary to establish a *bona fide* antigen. If these candidate autoantigens are validated, they could be incorporated into multi-analyte autoantigen panels for diagnostic or prognostic purposes.

Although many known antibody epitopes contain a linear or contiguous segement, those with purely conformational epitopes or mimotopes may not be identified using PIWAS. PIWAS as presented, is limited to identifying linear epitopes at a proteome scale. Thus, we are developing PIWAS with degenerate positions that leverage motif patterns identified by IMUNE [24]. Furthermore, the current method uses the maximum signal observed within the protein sequence for a particular patient, but some antigens have multiple antibody epitopes [40]. The use of multiple signals within a protein is another avenue of development to improve both sensitivity and specificity of PIWAS.

In conclusion, we developed PIWAS to enable robust, proteome-wide, cohort-based antigen discovery. PIWAS analyzes the datasets resulting from random peptide library selections against case and control cohorts (e.g., SERA) to discover shared candidate antigens, regardless of whether the epitopes therein are public or private. Since SERA employs random libraries, PIWAS can be applied to multiple proteomes utilizing the same physical assay. As the size of case and control datasets continue to increase, PIWAS may uncover previously undiscovered antigens with potential utility in diagnostic and therapeutic applications. Finally, PIWAS may be useful to investigate, in an unbiased manner, the association of autoantigens, human pathogens, and commensal organisms with human disease.

## Materials and Methods

### Serum epitope repertoire analysis (SERA)

Development and preparation of the *Escherichia coli* random 12-mer peptide display library (diversity 8×10^9^) has been described previously [24]. SERA was performed as described [24]. Briefly, serum was diluted 1:25 and incubated for 1 hr with a 10-fold oversampling of the library (8×10^10^ cells/well) in a 96-well plate format at 4°C with orbital shaking (800 rpm) during which time serum antibodies bind to peptides on the bacterial surface that mimic their cognate antigens. Cells were then collected by centrifugation (3500 rcf × 7 min), the supernatant was removed, and the cell pellets were washed by resuspending in 750 µL PBS + 0.05% Tween-20 (PBST). The cells were again collected by centrifugation (3500 rcf × 7 min) and the supernatant was removed. Cell pellets were resuspended in 750 µL PBS and mixed thoroughly with 50 µL Protein A/G Sera-Mag SpeedBeads (GE Life Sciences, 17152104010350) (6.25 % the beads’ stock concentration). The plate was incubated for one hour at 4°C with orbital shaking (800 rpm). Bead-bound cells were captured in the plate using a Magnum FLX 96-ring magnet (Alpaqua, A000400) until all beads were separated. Unbound cells in the supernatant were removed by gentle pipetting, leaving only those cells bound to A/G beads. Beads were washed 5X by removing from the magnet, resuspending in 750 µL PBST, and then returning to the magnet. The supernatant was removed by gentle pipetting after the beads were securely captured. Cells were resuspended in 750 µL LB with 34 µg/mL chloramphenicol and 0.2% wt/vol glucose directly in the 96-deep-well plate and grown overnight with shaking (300 rpm) at 37°C.

### Amplicon library preparation for sequencing

After growth, cells were collected by centrifugation (3500 rcf for 10 min) and the supernatant was discarded. Plasmids encoding the selected peptides were isolated in 96-well format using the Montage Plasmid Miniprep_HTS_ Kit (MilliPore, LSKP09604) on a Multiscreen_HTS_ Vacuum Manifold (MilliPore, MSVMHTS00) following the “Plasmid DNA—Full Lysate” protocol in the product literature. For amplicon preparation, two rounds of PCR were employed; the first round amplifies the variable “X12” peptide region of the plasmid DNA. The second round barcodes each patient amplicon library with sample-specific indexing primers for data demultiplexing after sequencing. KAPA HiFi HotStart ReadyMix (KAPA Biosystems, KK2612) was used as the polymerase master mix for all PCR steps. Plasmids (2.5 µL/well) were used as template for a first round PCR with 12.5 µL of KAPA ReadyMix and 5 µL each of 1 uM forward and reverse primers. The primers (Integrated DNA Technologies) contain annealing regions that flank the X12 sequence (indicated in bold) and adapter regions specific to the Illumina index primers used in the second round PCR. Forward primer: TCGTCGGCAGCGTCAGATGTGTATAAGAGACAGVBHDV**CCAGTCTGGCCAGGG** Reverse primer: GTCTCGTGGGCTCGGAGATGTGTATAAGAGACAG**GTGATGCCGTAGTACTGG** A series of five degenerate bases in the forward primer, VBHDV (following IUPAC codes), provide base diversity for the first five reads of the sequencing on the NextSeq platform. The five base pairs were designed to be non-complementary to the template to avoid bias during primer annealing. To reduce non-specific products, a touchdown PCR protocol was used with an initial annealing temperature of 72°C with a decrease of 0.5°C per cycle for 14 cycles, followed by 10 cycles with annealing at 65°C. The 25 uL primary PCR product was purified using 30uL Mag-Bind TotalPure NGS Beads (Omega Bio-Tek, M1378-02) according to the manufacturer’s protocol. The second round PCR (8 cycles, 70°C annealing temperature) was performed using Nextera XT index primers (Illumina, FC-131-2001) which introduce 8 base pair indices on the 5’ and 3’ termini of the amplicon for data demultiplexing of each sample screened. The PCR 1 product (5uL) was used as a template for the second PCR with 5uL each of forward and reverse indexing primers, 5uL PCR grade water and 25uL of KAPA ReadyMix. The PCR product (50uL) was cleaned up with 56 uL Omega Mag-Bind TotalPure NGS Beads per reaction. A 96-well quantitation was performed using the Qubit dsDNA High Sensitivity assay (Invitrogen, Q32851) adapted for a microplate fluorimeter (Tecan SPECTRAFlour Plus) measuring fluorescence excitation at 485 nm and emission at 535 nm. Positive (100 ng) and negative (0 ng) controls, included with the Qubit kit, were added to the plate as standards along with 2uL of each PCR product diluted 1:100 for quantitation. The fluorescence data were used to calculate DNA concentration in each well based on the kit standards. To normalize the DNA and achieve equal loading of each patient sample on NGS, the DNA in each well was diluted with Tris HCl (pH 8.5, 10 mM) to 4 nM and an equal volume from each well was pooled in a Lo-Bind DNA tube for sequencing.

The sample pool was prepared for sequencing according to specifications of the Illumina NextSeq 500. Due to the low diversity in the adapter regions of the amplicon after the first five bases, PhiX Run Control (Illumina, FC-110-3001) was included at 40% of the final DNA pool. The pool was sequenced using a High Output v2, 75 cycle kit (Illumina, FC-404-2005).

### Naïve Library Sequencing

An aliquot of the naïve X12 library representing 10-fold oversampling of the diversity was divided into 10 tubes, and the plasmids were purified and amplicons prepared as described above. Each prep was barcoded with a unique set of indices and sequenced on the NextSeq 500 to yield approximately 400 million unique sequences.

### Cohorts

#### Control cohort

Specimens from 1,157 apparently healthy individuals were used as a control cohort.

#### SLE cohort

De-identified specimens from 31 individuals diagnosed with SLE, and primarily female (27), were acquired from Proteogenex (9) and BioIVT (22). The mean age within this cohort was 43 years, with a range of 22-72.

#### Anti-Smith cohort

Samples from 34 subjects that tested positive for Anti-SM RNP (4) or Anti-Smith (30) antibodies by predicate ANA multiplex testing were obtained from Discovery Life Sciences. Subjects ranged in age from 18 -74, with the majority (26) being female.

### PIWAS Calculation

We define case (*T*), and control (*U*), cohorts of samples and begin with 12mer amino acid sequences for each sample generated by SERA (minimum of 1e6 total unique sequences per sample).

#### Enrichment calculation

We decompose each 12 mer from SERA into constituent *k*mers (where *k*=5 and *k*=6 consecutive amino acids). For every *kmer* in each sample (*S*), we calculate enrichment as:

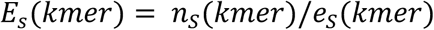

where *n(kmer)* is the number of unique 12mers containing a particular *kmer* and *e*_*s*_(*kmer*) is the expected number of *kmer* reads for the sample, defined as:

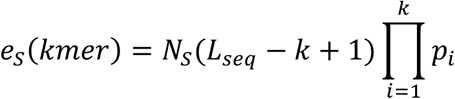

where *N*_*s*_ is the number of 12mer reads generated for *S, L*_*seq*_is the length of the amino acid reads (12), *k* is the kmer length, and *p*_*i*_ is the amino acid proportion for the *i*th amino acid in *kmer* in all 12mers from *S*.

#### Number of standard deviation normalization

For every kmer, we normalize enrichment values to a control population. We define the control enrichment values as:

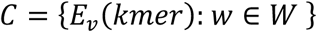

where *W* is the control cohort (*U*).

The normalized enrichment is calculated as:

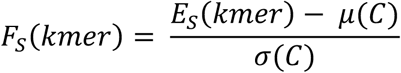

where *μ*(*C*) is the mean of C and σ(*C*) is the standard deviation of C.

#### PIWAS score calculation

For each protein *p* and sample *s*, we calculate a PIWAS score *P(s,p)*, defined as:

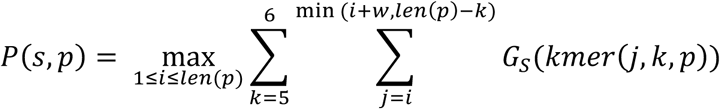

where *w* is the width of the smoothing window, *len(p)* is the length of protein *p, kmer(j,k,p)* is the kmer of length *k* at location *j* in protein *p*, and *G*_*s*_ is either *E*_*s*_ or *F*_*s*_. Similarly, we record the location of this maximum statistics value, *P*_*loc*_(*s, p*), as:

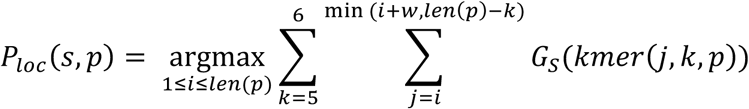

#### Cohort comparison statistics

For each protein *p*, we define our case enrichments as:

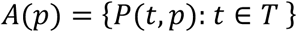

Similarly, we define our control enrichments as:

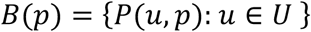

We use several statistical tests to compare *A(p)* and *B(p)*, including traditional tests like the Mann-Whitney U and Kolmogorov-Smirnov. We calculate effect size as the Hedges’ *g* statistic. We calculate the Outlier Sum, which we define as *O(p)*, statistic defined in Tibshirani and Hastie [38]. We perform 1,000 random permutations of the samples in *A(p)* and *B(p)* and calculate the Outlier Sum to calculate *0*^0^(*p*), the null distribution of the Outlier Sum for protein *p*. We calculate the z-score as:

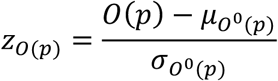

Since the Outlier Sum is a sum of i.i.d. variables, we can apply the Central Limit Theorem and calculate a p-value for *z*_*O*(*p*)_ using the normal distribution.

We define the sets of case and control locations as:

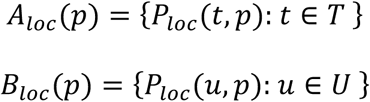

We perform a Kolmogorov-Smirnov test comparing *A*_*loc*_ (*p*) and *B*_*loc*_ (*p*) to identify proteins with locational conservation of epitopes.

### Proteome description

The reference *Homo sapiens* proteome was downloaded from Uniprot[41] on February 28, 2019.

### Kmer Enrichment Analysis

We compared the count of unique kmer species vs. enrichment scores for 5 and 6 mers in assays with a random library vs. those incubated with serum. We also compared the distribution of PIWAS values and average PIWAS values across control and SLE samples.

### Autoantigen Simulation Experiments

To simulate the effects of changing the magnitude and prevalence of autoantigenic signal, the real PIWAS signal against one of the Smith antigens in the SLE cohort was selected for use in a series of simulations (P14678: Small nuclear ribonucleoprotein-associated proteins B and B’). For every sample, the PIWAS values were calculated. To simulate different magnitudes of effect, the SLE PIWAS values were multiplied by scaling factors ranging from [0.1,2] and the outlier sum statistics were calculated relative to unscaled control values. To simulate different prevalences of effect, the SLE PIWAS values were divided into “high” (PIWAS > 6) and “low”(PIWAS < 6) values, 1000 random samplings with replacement of the SLE cohort were taken to simulate prevalences of “high” ranging from [0.01, 1], and the outlier sum statistics were calculated relative to unaffected control values.

### Data Availability

PIWAS scores for the the human proteome in the SLE, anti-Smith, and control samples have been provided as a supplemental file.

## Author contributions

Conceptualization: WH, KK, PD, JS; Data curation: WH; Formal analysis: WH; Funding acquisition: PD, JS; Software: WH; Visualization: WH; Writing-Review & Editing: WH, KK, PD, JS; Writing-Original draft preparation: WH, JS.

## Acknowledgements

We thank the Serimmune team for supporting the development of PIWAS and processing of samples: Becky Waitz, Jack Reifert, Joel Bozekowski, Burak Himmetoglu, Brian Martinez, Gregory Jordan, Timothy Johnston, Cameron Gable, Steve Kujawa, Elisabeth Baum-Jones. Special thanks to Elizabeth Stewart for editing this manuscript.

